# DHX9-dependent recruitment of BRCA1 to RNA is required to promote DNA end resection in homologous recombination

**DOI:** 10.1101/2019.12.20.884593

**Authors:** Prasun Chakraborty, Kevin Hiom

**Affiliations:** Division of Cellular Medicine, Jacqui Wood Cancer Centre, School of Medicine, University of Dundee. DD1 9SY. Scotland. United Kingdom

## Abstract

DExD/H-box helicase 9 (DHX9/RNA helicaseA) is ubiquitously expressed ATP-dependent helicase that unwinds a variety of complex DNA and RNA secondary structures, suggesting it might have a role in DNA replication and the repair of DNA damage. Here we identify a pivotal role for DHX9 in homologous recombination (HR). Cells that are deficient in DHX9 are impaired in the generation of single stranded DNA (ssDNA) by DNA end resection. Consequently these cells fail to recruit RPA and RAD51 to sites of DNA damage and are hypersensitive to treatment with the DNA damaging agents camptothecin and Olaparib. A critical early step in the initiation of HR is the recruitment of BRCA1 mediator protein to DNA damage to promote DNA resection. We show here that recruitment of BRCA1 to DNA damage foci is dependent on its interaction with DHX9 to form the BRCA1-D complex, which binds to nascent RNA. Together our data identify a pivotal role for DHX9 in homologous recombination and highlight the important contribution of RNA in the recruitment of BRCA1 to DNA damage for the repair of DNA breaks

## Introduction

DHX9 is a multifunctional DNA/RNA helicase that has been implicated in a variety of cellular functions that are primarily linked to the transcription and processing of RNA(1). In cells, DHX9 promotes the assembly of RNA binding proteins on newly transcribed RNA(2) and resolves large RNA secondary structures that arise after transcription of genomic inverted Alu repeats(3). In mice, homozygous deletion of DHX9 results in embryonic lethality(4) and in human diploid fibroblasts, suppression of DHX9 leads to blocked replication, P53-mediated arrested growth and senescence(5), indicating that it performs an essential function. Conversely, upregulation of DHX9 is required for tumour growth in multiple tumour types.

DHX9 is an SF2 type DExH-box helicase, which has two binding sites for double-stranded RNA in its N-terminal region and a glycine-rich RGG region that enables it to bind single-stranded nucleic acid(1). Biochemical studies, *in vitro*, have shown that DHX9 unwinds DNA and RNA substrates with a 3’ to 5’ polarity, with a preference for RNA that has a 3’ single-stranded tail(6). It also unwinds a variety of complex nucleic acid secondary structures including RNA-DNA hybrid, D-loops, DNA forks and RNA and DNA guanine quadruplexes (G4-DNA/RNA)(6,7). In cells, DHX9 localizes to origins of replication and its suppression causes replicative stress leading to the hypothesis that it may play a role in DNA replication(1). DHX9 also exhibits functional similarities with the BLM and WRN helicases(8) and interacts with several DNA repair proteins, including BRCA1(9) and Ku86(10), suggesting that it could also play a role in the maintenance of genomic stability. However, a defined role for DHX9 in the processing and repair of DNA damage has not yet been demonstrated.

Repair of double stranded DNA breaks (dsb) is carried out by two major pathways, homologous recombination and non-homologous end joining (NHEJ). Whereas the former is operative during S and G2 phases the latter functions throughout the cell cycle. Both pathways contribute to the repair of canonical dsb, for example caused by exposure to IR or the action of endonucleases. However, breaks resulting from collapsed replication forks, such as those caused by the topoisomerase inhibitor camptothecin, are one-ended and require HR. In the absence of HR these breaks may be ligated to unrelated DNA ends through a long-range end-joining pathway mediated by 53BP1 causing the generation of potentially harmful translocations(11,12).

An increasing body of evidence suggests that RNA and RNA processing proteins play a role in the repair of DNA breaks(13). DNA damage response small non-coding RNAs (DDRNA) and damage-induced long non-coding RNAs (dilncRNA), which are transcribed in the vicinity of DNA breaks, promote the recruitment of DNA repair factors to DNA damage and may also bridge broken DNA ends to facilitate their repair. Some of these RNAs require processing by DROSHER and DICER indicating a potentially important role for the RNA interference pathway in the repair of DNA breaks(14,15). Although, RNA and RNA processing proteins influence the dsb-break repair, a consensus mechanism through which this occurs is yet to emerge. For example, while several RNA helicases have been found to associate with DNA damage it is unclear how RNA unwinding promotes the repair of dsb.

Here, we show that the RNA helicase DHX9 is required in the repair of DNA damage by the HR pathway. DHX9 accumulates at sites of DNA damage where its unwinding activity is required to promote the resection of broken DNA ends and binding of the single strand binding complex RPA. Moreover, DHX9 plays a critical role in the recruitment of BRCA1 to DNA damage through a mechanism that links the repair of dsb to newly transcribed RNA.

## Results

### DHX9 is redistributed in response to DNA damage

To determine if DHX9 is a component of the DDR we analysed its distribution in cells before and after treatment with DNA damaging agents. Cells were treated with camptothecin or ionizing radiation (IR) and then fixed and extracted to remove proteins that were not associated with chromatin. Cells were stained with antibody against DHX9 and its cellular distribution was visualized by fluorescence imaging. Treatment of cells with camptothecin caused DHX9 to accumulate in discreet nuclear foci (Figure 1A). These foci colocalized significantly with the phosphorylated form of histone H2AX (γH2AX), confirming that DHX9 is recruited to sites of DNA breaks (Figure 1B,C). However, unlike the phosphorylation of H2AX, which occurs within 30 minutes(16,17), the accumulation of DHX9 in nuclear foci was detected approximately 2 hours after exposure to DNA damage. This suggested that recruitment of DHX9 to DNA lesions might be a late event in the DDR, occurring after the initial detection and processing of DNA damage. Alternatively, DHX9 might only accumulate at sites of persistent DNA damage, which are difficult to repair. Although DHX9 foci were also detected in cells treated with IR, they did not significantly co-localize with γH2AX (Figure 1D). This might reflect a qualitative difference in the DNA breaks generated by IR and camptothecin, but could also be explained if DHX9 also forms foci at lesions other than DNA breaks.

**Figure 1.**
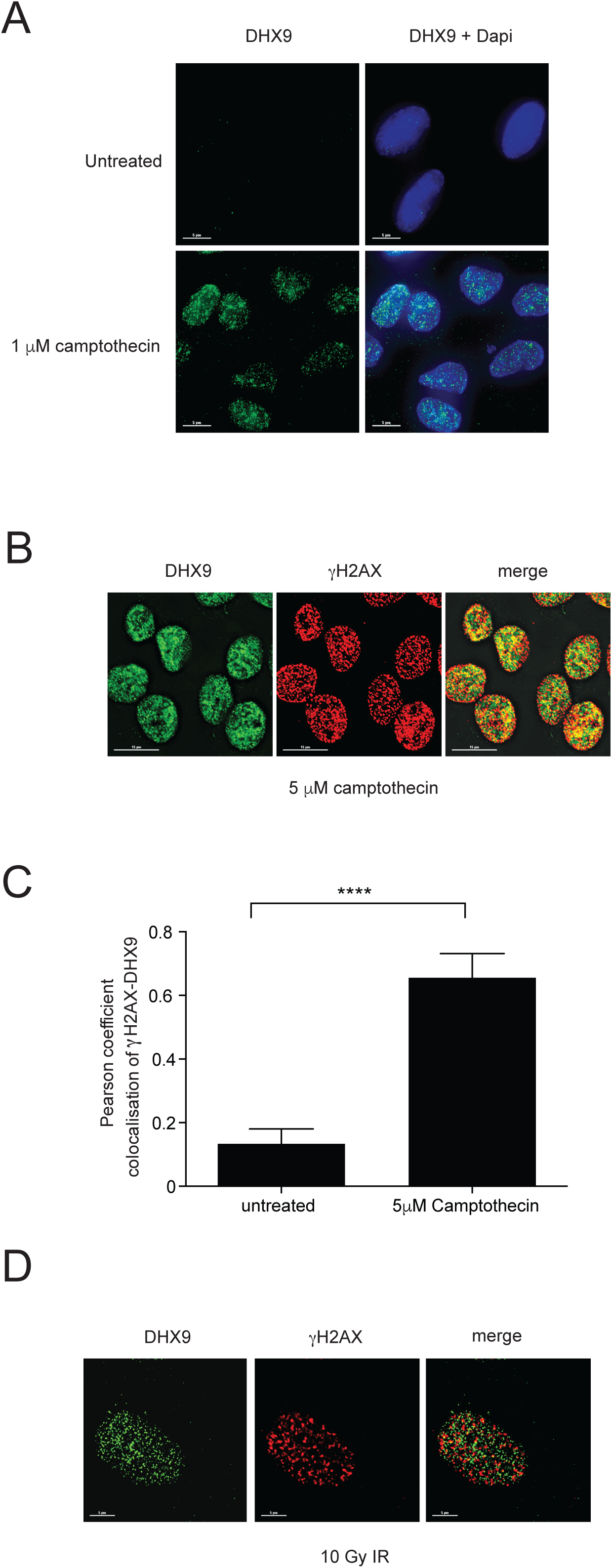
(**A**) Fluorescence image showing that DHX9 localizes into DNA damage nuclear foci in cells treated with camptothecin for 2 hours. Nuclei are stained with Dapi. (**B**) Fluorescence images showing that DHX9 (green) co-localizes with γH2AX (red) in cells treated with camptothecin for 2 hours. Co-localized proteins identified as yellow staining in the merged image. (**C**) Quantification of (**B**) showing pearson coefficient for co-localization of DHX9 and γH2AX in cells treated with camptothecin. Statistical significance was determined using one way Anova test (\*\*\*\**p* < 0.0001). (**D**) Fluorescence images showing that DHX9 (green) does not co-localize with γH2AX (red) in cells treated with Ionizing radiation.

*In vitro*, DHX9 exhibits a preference for unwinding RNA substrates and in cells it associates with nascent RNA during transcription(2,6). It was interesting, therefore, that DNA damage induced DHX9 foci were greatly diminished in cells treated with RNaseA to degrade RNA (Supplementary Figure S1A-D). DHX9 foci were not affected in cells treated with RNaseH1 to degrade RNA-DNA hybrid. We concluded that the dynamic accumulation of DHX9 into foci in response to DNA damage is dependent, at least in part, on RNA.

### DHX9 is required for the homology dependent repair of DNA breaks

We next determined if DHX9 contributes to the repair of DNA damage. We used two different siRNAs to deplete DHX9 in U2OS cells and measured their sensitivity to different DNA damaging agents (Figure 2A). DHX9-defective cells were hypersensitive to treatment with camptothecin and etoposide, which inhibit Topisomerase I and topoisomerase II, respectively. They were also hypersensitive to Olaparib, an inhibitor of Poly-ADP ribose polymerase 1 (PARP1) (Figure 2B-D). By contrast, DHX9-defective cells were not sensitive to DNA damage caused by exposure to IR (Figure 2E).

**Figure 2.**
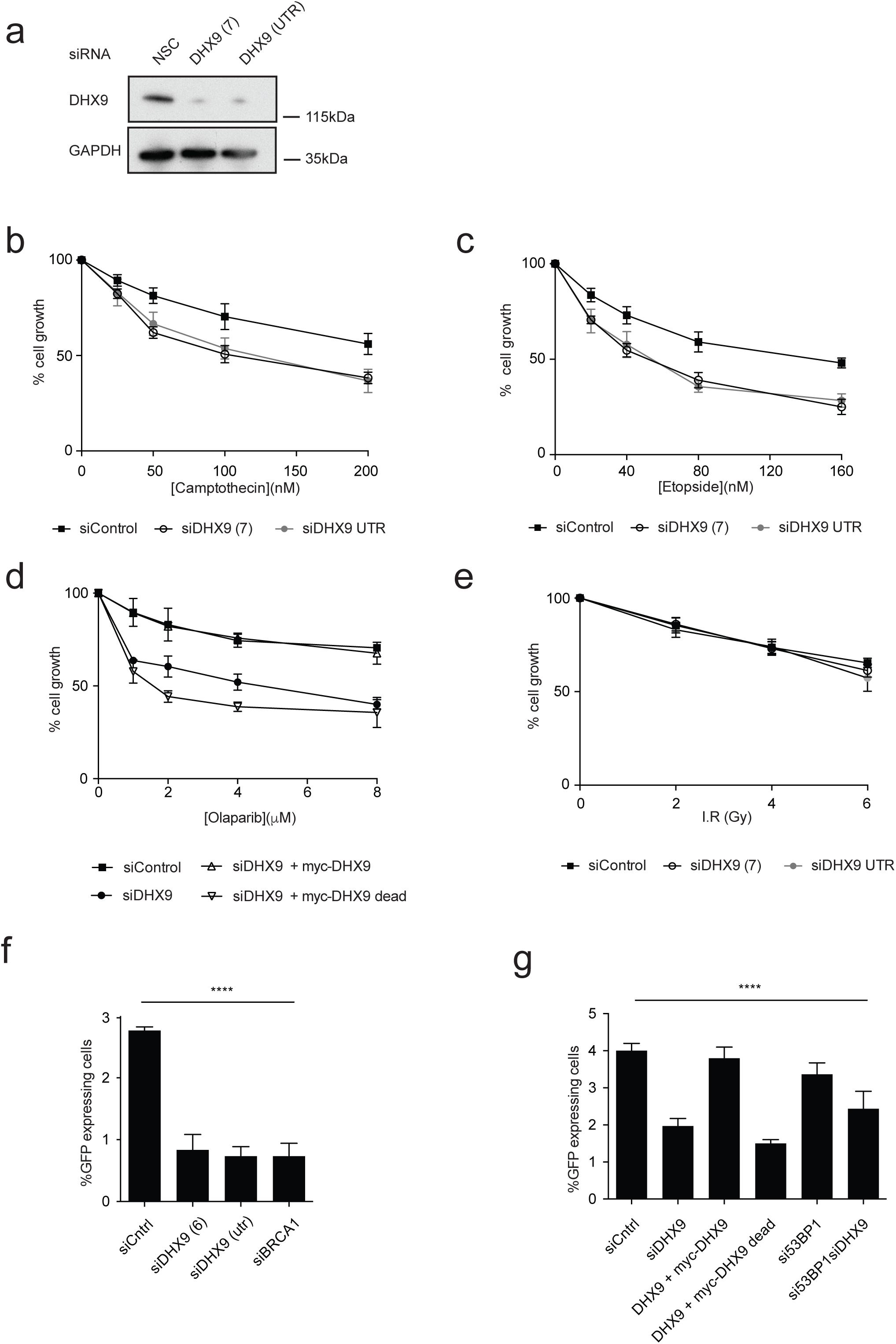
(**A**) Western blot showing knockdown by two different siRNAs. GAPDH was probed as a loading control. (**B**) Cells knocked down for DHX9 are hypersensitive to DNA damage caused by treating cells with camptothecin, (**C**) etoposide and (**D**) Olaparib. (**E**) DHX9 defective cells are not sensitive to treatment with ionizing radiation.(**F**) DHX9 deficient cells are defective in the repair of an I-Sce1 induced dsb by homologous recombination (HR) using a pDR-GFP reporter assay. Cells knocked down for BRCA1 are also indicated. Percentage GFP- expressing cells is shown. (**G**) HR-mediated dsb repair requires the helicase activity of DHX9. Over-expression of siRNA resistant myc-tagged wild type DHX9 but not helicase ‘dead’ D990A mutant DHX9 restores HR mediated repair and (**C**) Olaparib resistance in DHX9 knockdown cells. Statistical significance was determined using one way Anova test (\*\*\*\**p* < 0.0001)

Camptothecin, etoposide and Olaparib trap topisomerase I, topisomerase II and PARP1 on chromatin, respectively, causing stalling and collapse of replication forks and the generation of one-ended DNA breaks (oeb). Since oeb are commonly repaired by HR we hypothesized that DHX9 might play a role in HR mediated repair. Therefore, we used the established pDR-GFP cell reporter assay to measure HR-mediated repair of dsb induced by expression of I-Sce1 restriction enzyme(18). Cells knocked down for DHX9 exhibited a 50-70% reduction in HR-mediated repair of I-Sce1-induced dsb compared to wild type cells (Figure 2F). This was comparable with the HR defect measured for cells depleted of BRCA1, an important mediator protein for HR. Moreover, both HR-mediated dsb repair and resistance to Olaparib were restored in cells knocked down with an siRNA targeted to the 3’utr of DHX9 by overexpression of siRNA-resistant DHX9 cDNA from a plasmid, but not in cells expressing D990A DHX9 mutant protein that is specifically defective in its helicase activity (Figure 2D,G). This confirmed that the unwinding activity of DHX9 is critical for the repair of dsb by HR and prevents hypersensitivity to agents that introduce replication-blocking lesions into the genome.

### DHX9 promotes DNA end resection via its helicase activity

Repair of dsb by HR requires that broken DNA ends are first resected to generate regions of single stranded DNA (ssDNA), which are bound by single-strand binding protein complex RPA, protecting them from degradation. RPA is subsequently replaced on DNA by RAD51 recombinase, which catalyses DNA strand exchange and drives HR. To determine whether DHX9 is required in these early steps of the HR pathway we treated cells with IR and with camptothecin and used fluorescence imaging to detect γH2AX and RPA as a measure of DNA breaks and DNA end resection, respectively (Figure. 3A and B). Treatment with camptothecin induced similar levels of γH2AX foci in wild type cells and in cells depleted in DHX9, reflecting the generation of equivalent numbers of dsb (Figure 3A). However, cells that were depleted of DHX9 had many fewer RPA foci than did wild type cells, suggesting that DHX9 is required in the generation of ssDNA by DNA resection (Figure 3B, Supplementary Figure S1E). We then established that the generation of ssDNA required the helicase activity of DHX9 DNA by showing that RPA foci were restored to cells overexpressing wild type siRNA-resistant DHX9 cDNA from a plasmid but not the ‘helicase dead’ D990A mutant DHX9 (Figure 3C). Western blot analysis also confirmed that the level of RPA bound to chromatin was greatly reduced in cells depleted of DHX9 compared to control cells (Figure 3D). Finally, we used a BrdU staining assay to quantify ssDNA more directly, which confirmed that the generation of ssDNA was markedly reduced in DHX9 defective cells (Supplementary Figure S1E). Resection of DNA ends and binding of RPA are prerequisite for the recruitment of RAD51 to sites of DNA damage. Accordingly, cells knocked down in DHX9 were also greatly impaired in the formation of RAD51 foci in response to camptothecin-induced DNA damage (Figure 4A).

**Figure 3.**
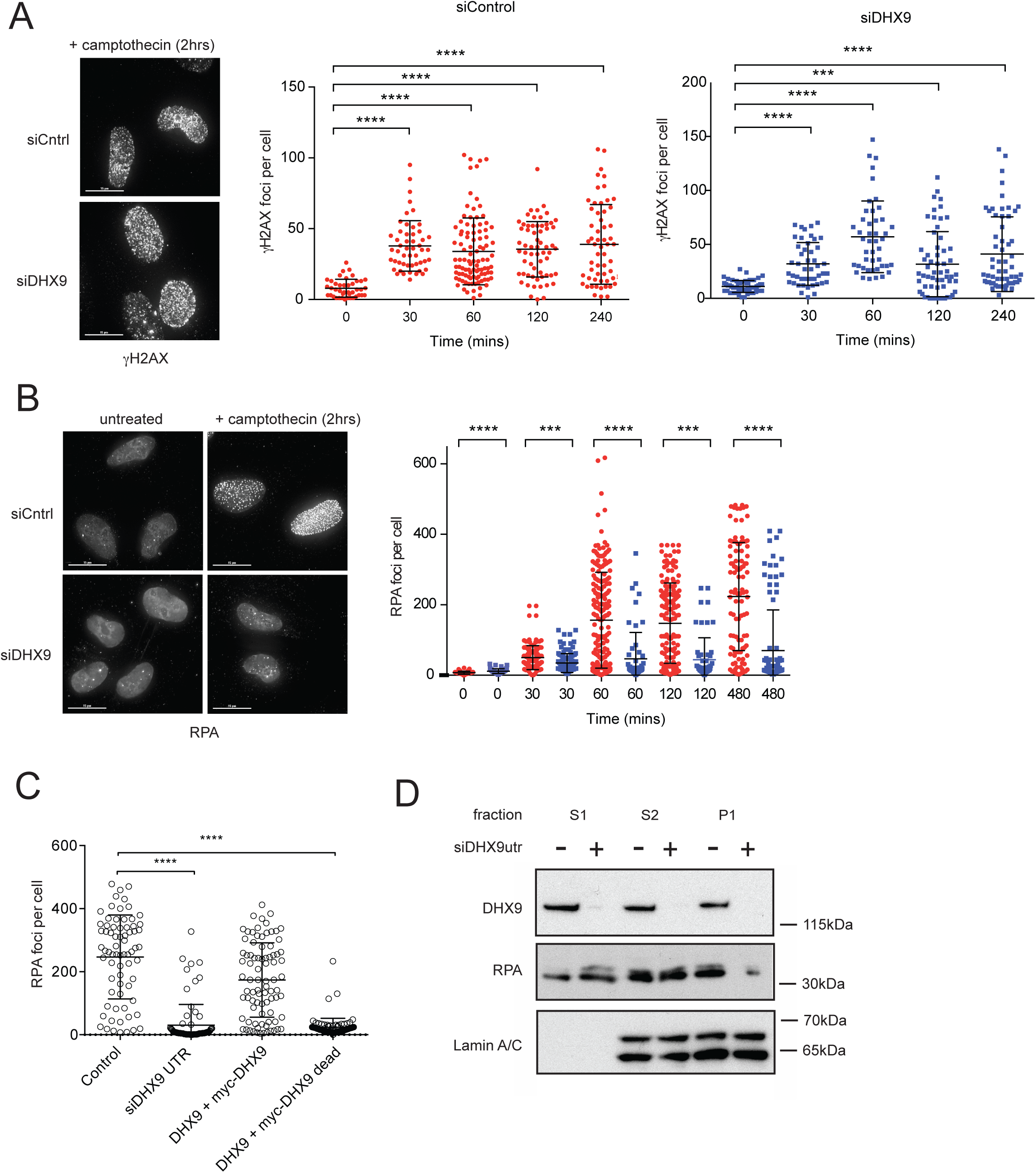
(**A**) Left panel-Fluorescent image showing similar levels of γH2AX foci in siControl and siDHX9 cells treated with 1 μM camptothecin for 2 hours. Right panel-Quantification of γH2AX foci over time (as indicated) in siControl (red) and siDHX9 (blue) cells. (**B**) Left panel- Fluorescent images showing that formation of RPA foci in siControl and siDHX9 cells treated with 1 μM camptothecin for 2 hours then recovered without drug for the indicated times. Right panel-Quantification showing impaired RPA foci in siDHX9 cells (blue) compared with siControl (red). (**C**) RPA foci are dependent on the helicase activity of DHX9 and are diminished in cells overexpressing siRNA resistant helicase dead D990A DHX9 mutant. Cells were treated with 2 μM camptothecin for 2 hours. Statistical significance of all graphs was determined using Mann–Whitney test (\*\*\*\**p* < 0.0001). (**D**) Western blot showing reduced RPA present in the chromatin (P1) fraction of cells knocked down for DHX9. Cytoplasmic (S1) and nucleoplasmic (S2) fractions are unchanged. Lamin A/C is shown as a control for nuclear fraction.

**Figure 4.**
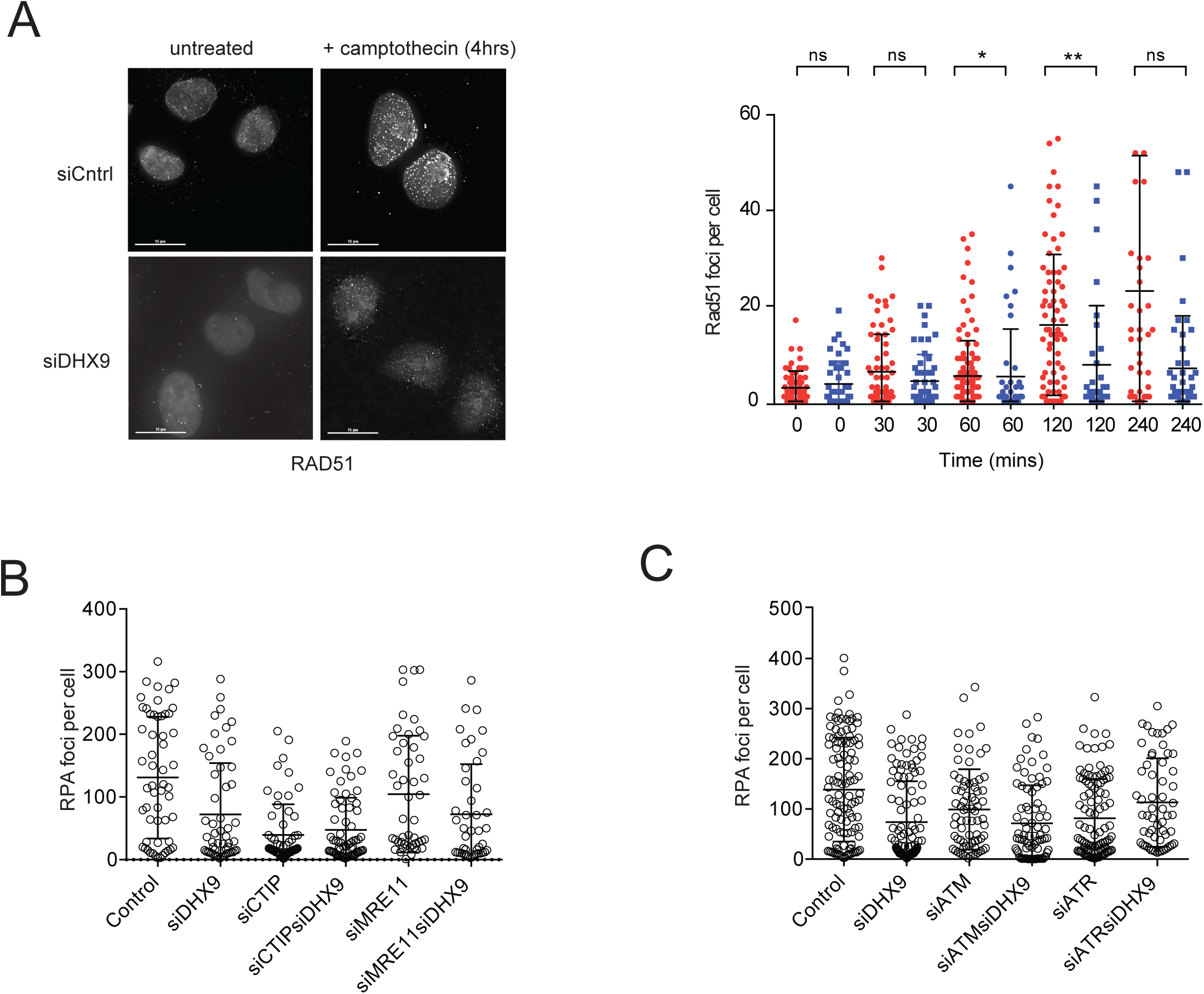
(**A**) Left panel-Fluorescent image showing decreased formation of RAD51 foci in cells knocked down for DHX9 treated with 1 μM camptothecin for 2 hours then left for 2 hours to recover. Right panel-Quantification of RAD51 foci over time in siControl (red) and siDHX9 (blue) cells. (**B**) DHX9 and CTIP are in a common genetic pathway for the formation of RPA foci. RPA foci in cells knocked down using siRNA for the indicated genes are shown. Cells were treated with 1 μM camptothecin for 2 hours. (**C**) DHX9, ATR and ATM are in a common genetic pathway for the formation of RPA foci. RPA foci in cells knocked down using siRNA for the indicated genes are shown. Statistical significance was determined using Mann–Whitney test (\*\*\*\**p* < 0.0001).

Although cells depleted of DHX9 were reduced in the recruitment of RPA and RAD51 to IR-induced DNA breaks (Supplementary Figure S2A-F), this defect was markedly smaller than that observed with camptothecin-induced damage. This was consistent with our earlier finding that colocalization of DHX9 with γH2AX is significantly less with IR-induced damage than with camptothecin damage. While it is possible that the pathways for resecting DNA damage generated by camptothecin and IR might be different in their requirement for DHX9, this difference might also reflect a greater contribution for NHEJ in the repair of IR-induced DNA breaks.

Resection of DNA ends is initiated by CTIP(19,20) and the MRN (MRE11-RAD50-NBS1) nuclease complex and is subsequently extended in the 3’ direction by EXO1 exonuclease, or through a combination of DNA2 and BLM (reviewed in(21)). To determine whether the contribution of DHX9 to the generation of ssDNA is through classical DNA end resection we looked at whether DHX9 is in the same genetic pathway as CTIP and MRE11. We confirmed that in wild type cells the majority of RPA foci generated in response to treatment with camptothecin were dependent on the activity of CTIP (Figure 4B). More than 50% of these foci were also dependent on DHX9. Since the mean number of RPA foci was not further diminished in cells depleted of both CTIP and DHX9, compared with cells depleted of CTIP alone, we concluded that DHX9 and CTIP function in a common pathway for the generation of ssDNA. Importantly, knockdown of DHX9 did not affect cellular levels of CTIP or BRCA1 as was reported for cells knocked down for the splicing factor SF3B1.

Mre11 was also required in the generation of RPA in response to camptothecin-induced DNA damage. However, its contribution to the formation of RPA foci was significantly less than that of either DHX9 or CTIP (Figure 4B). Nevertheless, because DNA resection was similar in cells depleted of both MRE11 and DHX9 as those impaired in DHX9 alone, we concluded that MRE11 and DHX9 also function in a common genetic pathway for the generation of ssDNA. Our data, suggest that DHX9 functions in the resection of DNA ends that is critical for the assembly of RAD51 nucleoprotein filaments to drive the HR pathway.

### DHX9 responds to signalling by ATR and ATM

We next addressed whether DHX9 responded to signalling by the DNA-damage signalling kinases ATM and ATR. In cells treated with camptothecin, knockdown of ATR and knockdown of DHX9 caused a similar decrease in RPA foci. Since knockdown of DHX9 and ATR together caused no further reduction in RPA foci we inferred that DHX9 mediated end resection responds to the ATR signalling pathway (Figure 4C). Knockdown of ATM in U2OS cells also impaired the generation of RPA foci in response to camptothecin-induced DNA damage. However, this defect was less severe than that caused by depletion of either DHX9 or ATR, confirming that ATR plays a bigger role than ATM in promoting DNA end resection in response to camptothecin-induced DNA damage. Nevertheless, since knockdown of DHX9 was epistatic to ATM for the recruitment of RPA, we concluded that these proteins also function in a common pathway. Hence, our genetic analysis supports the hypothesis that DHX9 promotes resection in response to signals from both ATM and ATR, but that ATR-dependent signalling comprises the major pathway for signalling camptothecin-induced DNA damage.

### DHX9 interacts with BRCA1 in response to DNA damage

Repair of DNA breaks by HR is dependent on the key mediator protein BRCA1(22). Cells that are defective in BRCA1 are impaired in DNA resection and fail to recruit RAD51 to sites of DNA damage. Cells knocked down for both BRCA1 and DHX9 exhibited a similar defect in HR mediated repair of I-Sce1 induced DNA breaks to cells depleted for these proteins individually. Hence, DHX9 and BRCA1 operate in the same genetic pathway for HR (Figure 5A). Accordingly, we detected increased colocalization of DHX9 and BRCA1 in DNA damage foci induced by camptothecin (Figure 5B). Moreover, the accumulation of BRCA1 in DNA damage foci was abrogated in cells knocked down for DHX9 using siRNA, indicating that DHX9 is required for the localization of BRCA1 at sites of DNA damage (Figure 5C,D). Since expression of the helicase dead D990A mutant of DHX9 did not restore BRCA1 foci in these cells, we inferred that the unwinding activity of DHX9 is important for the localization of BRCA1 to DNA damage (Figure 5E).

**Figure 5.**
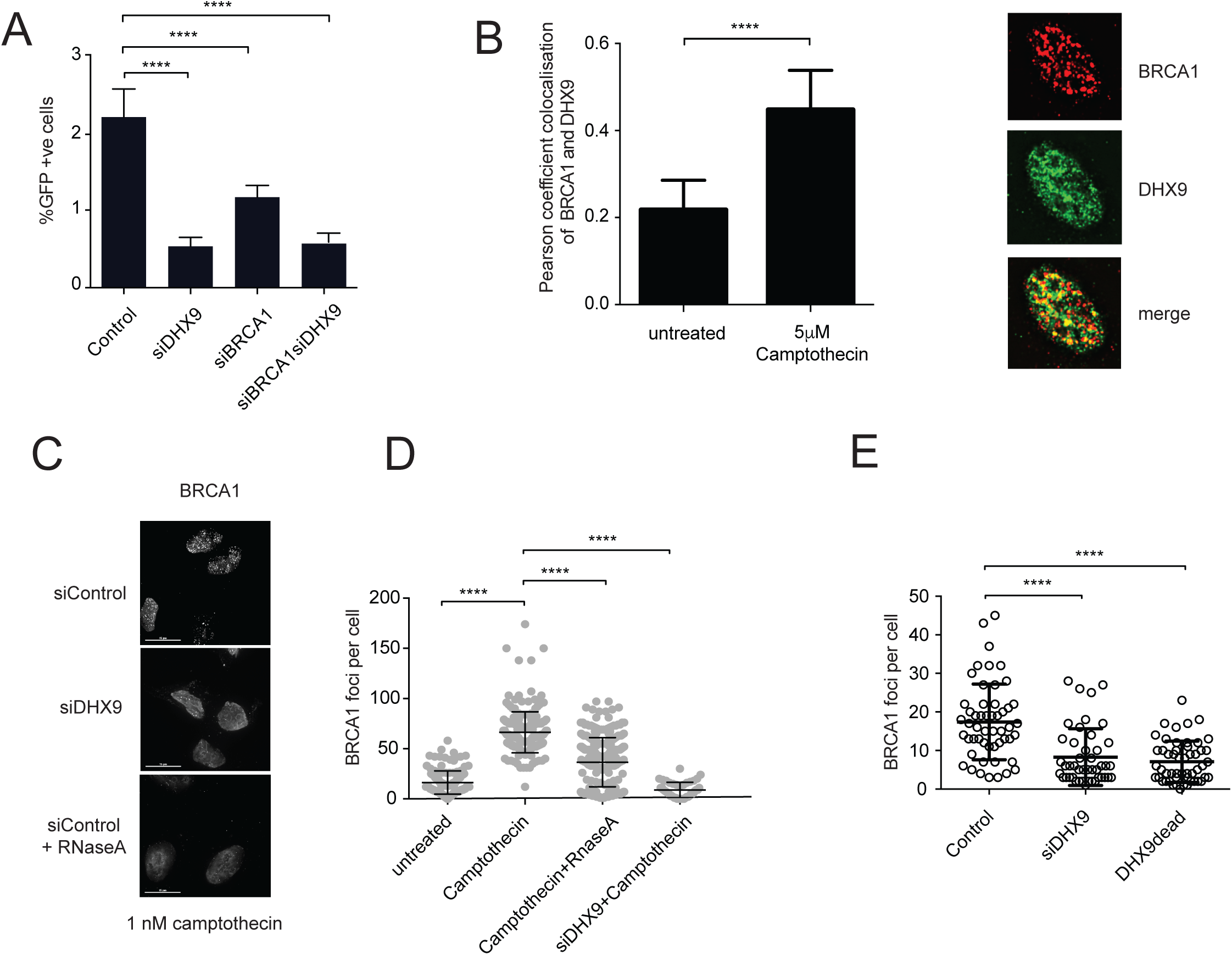
(**A**) BRCA1 and DHX9 are in the same genetic pathway for the repair of an I-Sce1 induced dsb by HR in a pDR-GFP assay. (**B**) BRCA1 and DHX9 co-localize in DNA Damage nuclear foci. Graph depicts pearson coefficient for co-localization of DHX9 and BRCA1 in untreated cells and cells treated with 1 nM camptothecin for two hours. (**C**) Fluorescent image showing that DNA damage induced BRCA1 foci are diminished in cells knocked down for DHX9 and in cells treated with RNaseA to degrade RNA. (**D**) Quantification of experiment in (**C**). (**E**) Graph showing that formation of BRCA1 DNA damage foci is dependent on the helicase activity of DHX9 and are not formed in cells expressing the helicase Dead D990A mutant DHX9 (indicated). Statistical significance was determined in A and B using one-way Anova test (\*\*\*\**p* < 0.0001) and in D and E using Mann–Whitney test (\*\*\*\**p* < 0.0001).

It was previously reported that DHX9 interacts with the c-terminal domain of BRCA1 in GST pull down experiments and in a yeast two hybrid assay(9,23). However, although very little endogenous BRCA1 co-immunoprecipitated with DHX9 in unperturbed cells (Figure 6A), their interaction was greatly enhanced in cells exposed to DNA damage (camptothecin, hydroxyurea and IR). Moreover, binding of BRCA1 to DHX9 was resistant to treatment with RNaseA indicating that is not mediated through RNA. We term this complex BRCA1-D (BRCA1-DHX9).

**Figure 6.**
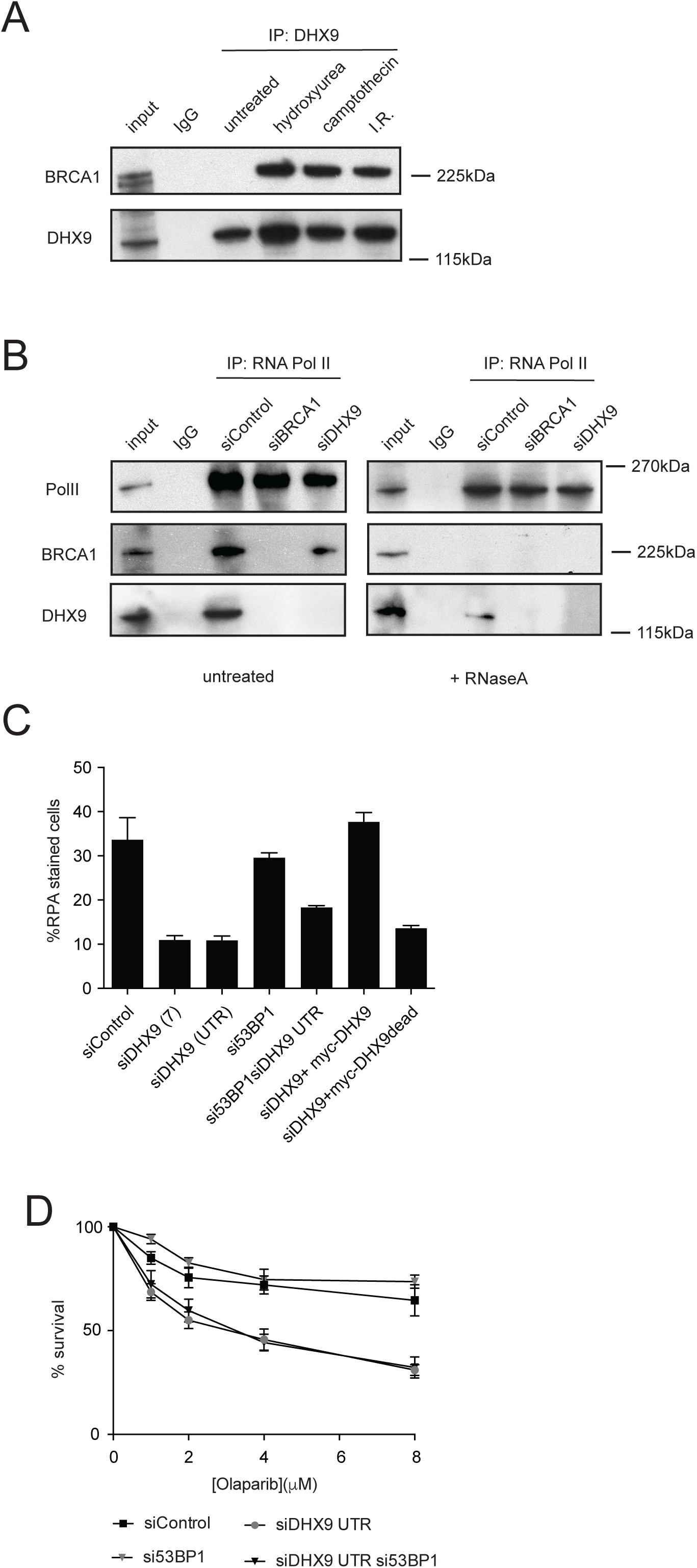
(**A**) Western blot showing co-immunoprecipitation of BRCA1 with DHX9 in cells treated with DNA damaging agents hydroxyurea (2 mM for 4 hours), camptothecin (2 μM for 2 hours) and ionizing radiation (10 Gy) (as indicated).(**B**) Western blot showing that BRCA1 and DHX9 co-purify with RNA Polymerase II that is dependent on RNA. Cells were treated with camptothecin (2 μM for 4 hours) before immunoprecipitation with antibody against RNA Pol II and probed with antibodies against RNA Pol II, DHX9 and BRCA1 as indicated. (**C**) Inhibition of 53BP1 restores RPA formation in DHX9 depleted cells. Percentage of cells stained for RPA measured using Fluorescent activated cell sorting. RPA is also restored by overexpression of DHX9 but not the helicase dead mutant DHX9 as indicated.(**D**) Knockdown of 53BP1 does not restore Olaparib resistance to DHX9 depleted cells. Statistical significance was determined using Mann–Whitney test (\*\*\*\**p* < 0.0001).

DHX9 has also been shown to bridge BRCA1 to the RNA Polymerase II holoenzyme(9). We recently showed that DHX9 recruits various proteins to RNA Pol II by facilitating their binding to nascent RNA(2). We confirmed that co-immunoprecipitation of BRCA1 with RNA Pol II is dependent on DHX9 and showed that this interaction was mediated by RNA (Figure 6B). While DHX9 and BRCA1 both co-immunoprecipitated with RNA Pol II, BRCA1 was greatly absent in samples that were treated with RNaseA and only a very small amount of DHX9 was retained. Furthermore, co-purification of BRCA1 with RNA Pol II was significantly reduced in cells that were depleted of DHX9. From this we infer that BRCA1 is recruited to the RNA Pol II transcription complex through its association with RNA and this recruitment requires DHX9.

In light of our biochemical analysis showing that DHX9-dependent recruitment of BRCA1 to RNA Poll II is dependent on RNA we examined whether the accumulation of BRCA1 in DNA damage induced foci was also dependent on RNA. Indeed, DNA damage induced BRCA1 foci were greatly diminished in cells treated with RNaseA, indicating that RNA is a component of these foci (Figure 5C,D). Together these observations suggest that a significant proportion of BRCA1 is recruited to chromatin by DHX9 as BRCA1-D and is mediated through its association with RNA. Furthermore, since BRCA1 plays a critical role in the initiation of DNA-end resection our data indicate that the impaired HR in cells depleted of DHX9 is explained by their failure to recruit BRCA1 to DNA damage.

### DHX9 contributes mechanistically to HR

Although BRCA1 is a key mediator of HR, it does not play an essential mechanistic role in the repair of dsb by this pathway. Instead it plays a critical role in channeling dsb for repair by HR and suppressing repair by 53BP1-mediated DNA end-joining in a process called pathway choice. This was established by showing that HR was restored in BRCA1-defective cells in which 53BP1 was mutated(11,24). Since DHX9 promotes the recruitment of BRCA1 in response to DNA damage we examined whether depletion of 53BP1 might also restore HR in cells that were knocked down for DHX9. However, this was not the case. In DHX9 depleted cells, neither, the defect in HR (Figure 2G), nor the defect in DNA resection (Figure 6C) was restored by the additional depletion of 53BP1. This was also true for hypersensitivity of DHX9 defective cells to PARP inhibitors (Figure 6D). Hence, we conclude that DHX9 does not promote HR simply by recruiting BRCA1 to sites of DNA damage to suppress 53BP1, but probably also contributes mechanistically to the repair of dsb by the HR pathway.

## Discussion

We have identified a critical role for DHX9 in the repair of DNA breaks by the HR pathway. We have demonstrated that DHX9 functions early in the HR process to promote the resection of broken DNA ends to ssDNA that is bound by RPA and subsequently by RAD51. Central to this function is the interaction of DHX9 with BRCA1 to form the BRCA1-D complex. We have shown that BRCA1-D is assembled in response to DNA damage, possibly through signalling by ATR/ATM kinases and is required for the recruitment of BRCA1 to sites of DNA damage. Consequently, cells lacking DHX9 are impaired in resection and in the repair of dsb by HR and are also hypersensitive to agents such as camptothecin and Olaparib, which generate replication blocking lesions.

Our data are consistent with a model in which BRCA1-D delivers BRCA1 to DNA damage where it can initiate DNA end resection. Previously, we showed that BRCA1 facilitates end resection by stimulating the activity of CTIP. This is consistent with our genetic data showing that DHX9 and CTIP operate in a common pathway for DNA resection that also requires DNA damage signalling by ATR/ATM. The initiation of HR for pathway choice, not only requires upregulation of DNA end resection but also the suppression of 53BP1-mediated end-joining by BRCA1. By recruiting BRCA1 to DNA damage, DHX9 is likely to be critical for pathway choice. However, since the HR defect in DHX9-depleted cells is not suppressed by knocking down 53BP1, we conclude that DHX9 must makes an additional mechanistic contribution to DNA end resection that is yet to be determined.

Interestingly, the contribution of DHX9 to DNA end resection is different depending on the type of DNA damage. While DHX9 co-localized with DNA breaks caused by camptothecin and Olaparib it did not co-localize with breaks introduced by treating cells with IR. Similarly DHX9-depleted cells were more profoundly defective in the resection of camptothecin induced DNA damage than that of IR. Accordingly, DHX9 defective cells are markedly more hypersensitive to treatment with camptothecin and Olaparib, than to IR. This might reflect a specific requirement for DHX9 in the processing of one-ended DNA breaks caused by collapsed replication forks that require repair by HR, compared with canonical two-ended DNA breaks, which are primarily repaired by NHEJ.

Our identification of DHX9 as a factor in DNA resection provides a direct link between HR mediated double strand break repair and RNA metabolism. DHX9 facilitates the assembly of RNA splicing factors on nascent RNA and we have shown here that DHX9 recruits BRCA1 to nascent RNA as part of the RNA Pol II transcription complex(2). Importantly these two proteins function as a bone fide complex since neither DHX9 nor BRCA1 bind to nascent RNA in the absence of the other. Moreover, since this complex is formed in response to DNA damage, it might serve as an important step in the regulation HR.

Intriguingly, recruitment of DHX9 and BRCA1 into DNA damage foci is also dependent on RNA. DNA damage foci formed by 53BP1 and MDC1 are also dependent on locally transcribed DDRNA, which do not play a role in the initial detection of DNA damage but are needed for secondary recruitment of DDR factors through a mechanism that is currently unknown(14,15). We have established here that nascent RNA is critical for the recruitment of BRCA1 to DNA damage and that this requires the assembly of the BRCA1-D complex as part of the DDR. Moreover, since DHX9 bridges the association of BRCA1 with RNA Pol II holoenzyme that the initiation of DNA end resection by BRCA1-D could be linked to slowing or paused transcription in regions of DNA damage(23). This is not unprecedented as BRCA1, together with another RNA helicase senataxin (SETX), have been shown to promote the repair of R-loop based DNA damage at transcriptional pause sites(25). It is possible, therefore, that BRCA1-D and BRCA1/SETX are manifestations of the same, or similar, transcription dependent repair activity.

Our work adds DHX9 to a growing number of RNA binding proteins that have been implicated in the maintenance of genome stability and the repair of DNA breaks. Importantly we have shown how DHX9 links RNA and RNA Pol II to the repair of DSB through the assembly of the BRCA1-D complex. Our demonstration that BRCA1-D, like BRCA1-A and BRCA1-C, plays a pivotal role in DNA end resection highlights the importance of this step in the regulation of dsb repair and uncovers a key contribution played by RNA in the repair of DNA damage and the maintenance of genomic stability(20,26-29).

## Methods

### Cell Culture

HeLa and U2OS cell lines were cultured in Dulbecco’s Modified Eagle’s Medium (DMEM) supplemented with 10 % fetal bovine serum (FBS) and 5 % ampicillin. All cell lines were obtained from ATCC.

### Antibodies, Chemicals and Reagents

Primary antibodies used in this study (concentrations used for IP and Western blot are indicated): rabbit polyclonal anti-RNA Helicase A (ab26271, Abcam, for IP 5 µg/ml, Immunofluorescence 1:1000 and WB 1:2000 dilution), mouse monoclonal anti-BRCA1 antibody (OP92, Ab-1; Calbiochem, for IPs 10μg/ml, IF 1:250 and WB 1:200 dilution), mouse anti-RPA32 (RPA2 Ab #2; Calbiochem, IF 1:500, WB 1:1000 and for FACS 1:100 dilution), rabbit polyclonal anti-RNA-Polymerase II (N-20; Santa Cruz Biotechnology, IP 4 μg/ml and WB 1:1000 dilution), mouse monoclonal anti-GAPDH antibody (GT239; GeneTex, WB 1:1000 dilution), mouse monoclonal anti-phospho-Histone H2A.X (clone JBW301; EMD Millipore, IF 1:250 dilution and WB 1:500 dilution), rabbit polyclonal anti-Rad51 antibody (H-92; Santa Cruz Biotechnology, IF 1:250 dilution), rabbit polyclonal anti-53BP1 Antibody (NB100-904; Novus Biologicals, IF 1:500, WB 1:2000), mouse monoclonal Anti-BrdU antibody (B44; BD Biosciences, 1:1000 dilution for IF). Secondary antibodies: goat anti-mouse Alexa Fluor 488 and 647 (Molecular Probes) was used at 1:200 dilution for FACS and 1:1000 dilution for IF, goat anti-rabbit Alexa fluor 488, 568 and 647 (Molecular Probes) was used at 1:1000 dilution for IF. For WB, Peroxidase-AffiniPure Goat Anti-Rabbit IgG (H+L) 111-035-144-JIR (1:5000 dilution) and Mouse IgG antibody (HRP) (GTX213111-01; GeneTex, 1:5000 dilution) were used. IgG controls were Normal Rabbit IgG #2729 (Cell Signaling) and normal mouse IgG (sc-2025; Santa Cruz Biotechnology). Hydroxyurea H8627 and (S)-(+)- Camptothecin C9911 (Sigma-Aldrich), AZD 2281-Olaparib (Axon Medchem), 5-Bromo-2’-deoxyuridine (Sigma, B5002), RNase A 19101 (17,500 U, Qiagen), Halt™ Protease Inhibitor Cocktail (100X) 78429 (Thermos Scientific) and PhosSTOP™ (Merck).

### Plasmids

pCMV-DHX9-GFPSPARK is designated pGFP-DHX9 and was purchased from Stratech (sino-biologicals). pGFP-DHX9dead contains D511A and E512A mutations made by site directed mutagenesis of pGFP-DHX9 using Q5 Site Directed Mutagenesis system (New England Biolabs) according to manufacturer’s instructions.

### SiRNA

The siRNAs were purchased from Dharmacon-ON-TARGET plus. Non-targeting siRNA Control-D-001810-01-05; DHX9-J-009950-07, DHX9utr-CTM-310164 and CTM-478066; 53BP1-J-003548-07; BRCA1-J-003461-09, ATR-L-003202-00, ATM-L-0030201-00, CTIP-J-011376-07, MRE11A-J-009271-07.

### Homologous recombination assay by FACS

75,000 cells were plated in 6-well plate overnight and transfected with siRNA. After 48 hours 2µg of the I-SceI expression vector pCBASce was transfected into cells. After a further 48 hours cells were harvested and the number of GFP-positive cells was measured by flow cytometry (Fortessa-BD Biosciences). For complementation experiments, plasmid expressing wild type of mutant DHX9 was transfected simultaneously with incubation of siRNA.

### Immunoprecipitation

Prior to immunoprecipitation, primary antibody was incubated with Dynabeads protein G beads (Invitrogen) overnight at 4 °C. Cells were lysed using Lysis Buffer (150 mM NaCl, 0.5% Triton X-100, 50 mM Tris-HCl pH 7.5, NaCl, 1 mM EDTA, 5% glycerol and 0.1% SDS) supplemented with protease and phosphatase inhibitor cocktails (Thermo Scientific). Lysed cells were passed through 23G needle 10 times on ice and incubated for 30 mins at 4 °C with shaking. For BRCA1, 500mM NaCl was included in the lysis buffer. Where indicated 3 μl of RNaseA was added to 1 ml of extract and incubated at 37 °C for 30 mins. Supernatant was cleared by high speed centrifugation at 14,000 rpm at 4 °C in a benchtop microcentrifuge. The cleared lysate (1 mg) was added to the antibody-Dynabead complex and incubated at 4 °C with rotation for between 4 and 12 hours. Immunocomplexes were separated using a magnet, washed three times in lysis buffer, boiled in sample buffer and loaded on a 4-12% Bis-Tris polyacrylamide gel (Invitrogen). Proteins were transferred to PVDF membrane using a Novex transfer system (Invitrogen) and immunoblotted using the indicated antibodies.

### Immunofluorescence microscopy

U2OS cells were seeded on coverslip overnight, non-chromatin bound proteins were pre-extracted using 0.5% TritonX-100 CSK buffer (25 mM Hepes pH 7.4, 50 mM NaCl, 1 mM EDTA, 3 mM MgCl_2_, 300 mM sucrose) for 2-5 mins on ice and then fixed with 3.7% paraformaldehyde. For RNase A treatment, cells were either treated for 30 mins at room temperature before or after fixation with RNase A (200 µg in 500μl PBS). Cells were washed three times in PBS and permeabilized for 10 mins in 0.5% Triton X-100/PBS. After three additional PBS washes, cells were blocked using 1-3% BSA/PBS for 30 mins. Cells were incubated with primary antibody (as indicated above), followed by the addition of Alexa Fluor-488 or −568 or 647 conjugated secondary antibodies (1:1,000). To visualize nuclei, cells were also stained with 0.5 μg/ml DAPI (Molecular Probes) for 15 mins. Slides were mounted using Prolong gold anti-fade reagent (Invitrogen) and images were acquired using a Delta vision DV4 wide-field deconvolution microscope with a 100X objective. Where indicated, cells were treated with 1 nM to 5 µM camptothecin and incubation for various times, after which cells were washed with PBS, pre-permeabilized, fixed and processed as above.

Cells were seeded on cover slips and grown overnight and then transfected with siRNA for 6 hours the media changed and then cells grown for a further 48 hours. Where indicated plasmid expressing wild type DHX9 or S990A mutant was transfected using lipofectamine 2000 reagent (Invitrogen) for 48 hours. Images were analyzed using a Delta Vision microscope and quantified using image J software.

Image Analysis-Colocalization between X protein and Y protein was assessed in single-cell regions using SoftWoRx (version 5.5.0, release 6). A rectangular selection was defined for each cell in deconvolved images and the Pearson coefficient of correlation was calculated for the volume. Nuclear foci counting macro. An ImageJ macro was used to count green and red foci inside nuclei. Nuclei were automatically detected in the DAPI channel using the Default ImageJ auto-thresholding method, and foci in the red and green channels were counted using ImageJ’s “Find Maxima” feature with a user-defined Prominence above background.

### ssDNA detection assay by Immunofluorescence

Cells were grown on microscope slides overnight. siRNA knockdown was performed as above and cells were incubated with 10 µM BrdU 24 hrs. 1 µM camptothecin was added in the media for 2 hours and then washed out with PBS. Cells were pre-permeabilized in ice for 5 mins then fixed and treated as above using non-denaturing conditions. Cells were incubated with primary antibody against BrdU overnight then washed and secondary antibody added as above. Images were captured with a Delta vision DV4 wide-field deconvolution microscope using a 40 × objective.

### Cell growth assay

U2OS cells were transfected with siRNA for 48 hours as above. 5,000 cells per well were seeded into a 12-well plate (time 0). At the indicated times, cells were recovered from the monolayer using trypsin and counted using Casey Cell Counter. Where indicated camptothecin and Olaparib were included in the medium and replenished every three days. Etoposide was added to medium for 12 hours and then replaced with fresh medium. Growth curves were plotted using data from three to six independent biological replicates.

### Flow Cytometry

Analysis of RPA staining by FACS was performed as described in (Jackson et al paper, Cytometry A. 2012 Oct; 81A(10): 922–928. Published online 2012 Aug 14. doi: 10.1002/cyto.a.22155) with a few modifications. Firstly, non-chromatin bound proteins were extracted with 0.5% Triton X-100 (PBS-T) for 15 min on ice followed by fixation in 4% para formaldehyde and then permeabilized for 30 mins at room temperature. After washing in PBS the cells were incubated with primary and secondary antibody subsequently, washed and stained with 0.02 % sodium azide, 250 μg/ml RNase A and 2 μg/ml of 4’,6-diamidino-2-phenylindole (DAPI) in PBS for at least 30 min at 4°C prior to analysis using a Fortessa flow cytometer (Beckton Dickinson). Data was collected from more than 3 independent experiments and quantified using Prism v6 (GraphPad Software).

### Cell Fractionation

Briefly, 3 × 10^6^ HeLa cells per condition were collected and suspended in 250 µl of buffer A (10 mM HEPES pH 7.8, 10 mM KCl, 1.5 mM MgCl_2_, 0.25 M sucrose, 10% glycerol, 1 mM DTT, 0.1% Triton-X-100, protease and phosphatase inhibitors) and incubated for 5 min on ice. The soluble cytoplasmic fraction (S1) was separated from the nuclei (P2) by centrifugation for 4 min at 1700 × *g* at 4°C. The nuclear fraction P2 was washed twice with 500 µl buffer A and suspended in 200 µl buffer B (3 mM EDTA, 0.2 mM EGTA, 1 mM DTT, phosphatase and protease inhibitors) and incubated at 4 °C for 30 min. The insoluble chromatin fraction (P3) was separated from nuclear soluble proteins (S3) pellet by acid-extraction using 0.25 N HCl and incubated on ice for 30 min. The lysate was then centrifuged at 16,000 x g for 15 min at 4°C. The supernatant (contains acid soluble proteins) was neutralized using 1 M Tris-HCl pH 8 using 1:5 volume. 25 µg of the protein was loaded in 4-12% Bis-Tris gradient polyacrylamide gel and transferred onto PVDF membrane (Millipore). Membranes were blocked in PBS with 0.1% Tween 20 and probed with respective primary antibodies. Bound proteins were detected with Pierce ECL western blotting substrates and developed with x-ray film (Konica Minolta).

### Data processing and statistical analysis

Values are shown with the standard deviation from at least three independent experiments unless indicated otherwise. Data were analyzed and, where appropriate, the significance of the differences between the mean values was determined using two-tailed Mann–Whitney unpaired t-test (**P≤0.01, ****P≤0.0001) or one-way Anova test as indicated. All statistics were performed using Prism v6 (GraphPad Software).

## Funding

This manuscript was funded by BBSRC [BB/P021387/1 to KH]; and by the Ninewells Cancer Campaign.

## Acknowledgements

We thank Professor Jerry Pelletier (McGill university, Quebec, Canada) for kind gift of plasmid expressing DHX9-Myc and Dr Graeme Ball (Dundee Imaging Facility, University of Dundee) for assistance with image analysis.

## References

1. Lee, T. and Pelletier, J. (2016) The biology of DHX9 and its potential as a therapeutic target. Oncotarget, 7, 42716–42739.

2. Chakraborty, P., Huang, J.T.J. and Hiom, K. (2018) DHX9 helicase promotes R-loop formation in cells with impaired RNA splicing. Nat Commun, 9, 4346.

3. Aktas, T., Avsar Ilik, I., Maticzka, D., Bhardwaj, V., Pessoa Rodrigues, C., Mittler, G., Manke, T., Backofen, R. and Akhtar, A. (2017) DHX9 suppresses RNA processing defects originating from the Alu invasion of the human genome. Nature, 544, 115–119.

4. Lee, C.G., Eki, T., Okumura, K., da Costa Soares, V. and Hurwitz, J. (1998) Molecular analysis of the cDNA and genomic DNA encoding mouse RNA helicase A. Genomics, 47, 365–371.

5. Lee, T., Di Paola, D., Malina, A., Mills, J.R., Kreps, A., Grosse, F., Tang, H., Zannis-Hadjopoulos, M., Larsson, O. and Pelletier, J. (2014) Suppression of the DHX9 helicase induces premature senescence in human diploid fibroblasts in a p53-dependent manner. J Biol Chem, 289, 22798–22814.

6. Chakraborty, P. and Grosse, F. (2011) Human DHX9 helicase preferentially unwinds RNA-containing displacement loops (R-loops) and G-quadruplexes. DNA Repair (Amst), 10, 654–665.

7. Chakraborty, P. and Grosse, F. (2010) WRN helicase unwinds Okazaki fragment-like hybrids in a reaction stimulated by the human DHX9 helicase. Nucleic Acids Res, 38, 4722–4730.

8. Brosh, R.M., Jr., Majumdar, A., Desai, S., Hickson, I.D., Bohr, V.A. and Seidman, M.M. (2001) Unwinding of a DNA triple helix by the Werner and Bloom syndrome helicases. J Biol Chem, 276, 3024–3030.

9. Nakajima, T., Uchida, C., Anderson, S.F., Lee, C.G., Hurwitz, J., Parvin, J.D. and Montminy, M. (1997) RNA helicase A mediates association of CBP with RNA polymerase II. Cell, 90, 1107–1112.

10. Rampakakis, E., Di Paola, D. and Zannis-Hadjopoulos, M. (2008) Ku is involved in cell growth, DNA replication and G1-S transition. J Cell Sci, 121, 590–600.

11. Bunting, S.F., Callen, E., Wong, N., Chen, H.T., Polato, F., Gunn, A., Bothmer, A., Feldhahn, N., Fernandez-Capetillo, O., Cao, L. et al. (2010) 53BP1 inhibits homologous recombination in Brca1-deficient cells by blocking resection of DNA breaks. Cell, 141, 243–254.

12. Bunting, S.F. and Nussenzweig, A. (2013) End-joining, translocations and cancer. Nat Rev Cancer, 13, 443–454.

13. Michelini, F., Jalihal, A.P., Francia, S., Meers, C., Neeb, Z.T., Rossiello, F., Gioia, U., Aguado, J., Jones-Weinert, C., Luke, B. et al. (2018) From “Cellular” RNA to “Smart” RNA: Multiple Roles of RNA in Genome Stability and Beyond. Chem Rev, 118, 4365–4403.

14. Francia, S., Cabrini, M., Matti, V., Oldani, A. and d’Adda di Fagagna, F. (2016) DICER, DROSHA and DNA damage response RNAs are necessary for the secondary recruitment of DNA damage response factors. J Cell Sci, 129, 1468–1476.

15. Francia, S., Michelini, F., Saxena, A., Tang, D., de Hoon, M., Anelli, V., Mione, M., Carninci, P. and d’Adda di Fagagna, F. (2012) Site-specific DICER and DROSHA RNA products control the DNA-damage response. Nature, 488, 231–235.

16. Rogakou, E.P., Boon, C., Redon, C. and Bonner, W.M. (1999) Megabase chromatin domains involved in DNA double-strand breaks in vivo. J Cell Biol, 146, 905–916.

17. Paull, T.T., Rogakou, E.P., Yamazaki, V., Kirchgessner, C.U., Gellert, M. and Bonner, W.M. (2000) A critical role for histone H2AX in recruitment of repair factors to nuclear foci after DNA damage. Curr Biol, 10, 886–895.

18. Pierce, A.J., Johnson, R.D., Thompson, L.H. and Jasin, M. (1999) XRCC3 promotes homology-directed repair of DNA damage in mammalian cells. Genes Dev, 13, 2633–2638.

19. Huertas, P. and Jackson, S.P. (2009) Human CtIP mediates cell cycle control of DNA end resection and double strand break repair. J Biol Chem, 284, 9558–9565.

20. Sartori, A.A., Lukas, C., Coates, J., Mistrik, M., Fu, S., Bartek, J., Baer, R., Lukas, J. and Jackson, S.P. (2007) Human CtIP promotes DNA end resection. Nature, 450, 509–514.

21. Cejka, P. (2015) DNA End Resection: Nucleases Team Up with the Right Partners to Initiate Homologous Recombination. J Biol Chem, 290, 22931–22938.

22. Moynahan, M.E., Chiu, J.W., Koller, B.H. and Jasin, M. (1999) Brca1 controls homology-directed DNA repair. Mol Cell, 4, 511–518.

23. Anderson, S.F., Schlegel, B.P., Nakajima, T., Wolpin, E.S. and Parvin, J.D. (1998) BRCA1 protein is linked to the RNA polymerase II holoenzyme complex via RNA helicase A. Nat Genet, 19, 254–256.

24. Bouwman, P., Aly, A., Escandell, J.M., Pieterse, M., Bartkova, J., van der Gulden, H., Hiddingh, S., Thanasoula, M., Kulkarni, A., Yang, Q. et al. (2010) 53BP1 loss rescues BRCA1 deficiency and is associated with triple-negative and BRCA-mutated breast cancers. Nat Struct Mol Biol, 17, 688–695.

25. Hatchi, E., Skourti-Stathaki, K., Ventz, S., Pinello, L., Yen, A., Kamieniarz-Gdula, K., Dimitrov, S., Pathania, S., McKinney, K.M., Eaton, M.L. et al. (2015) BRCA1 recruitment to transcriptional pause sites is required for R-loop-driven DNA damage repair. Mol Cell, 57, 636–647.

26. Coleman, K.A. and Greenberg, R.A. (2011) The BRCA1-RAP80 complex regulates DNA repair mechanism utilization by restricting end resection. J Biol Chem, 286, 13669–13680.

27. Wang, B., Matsuoka, S., Ballif, B.A., Zhang, D., Smogorzewska, A., Gygi, S.P. and Elledge, S.J. (2007) Abraxas and RAP80 form a BRCA1 protein complex required for the DNA damage response. Science, 316, 1194–1198.

28. Hu, Y., Scully, R., Sobhian, B., Xie, A., Shestakova, E. and Livingston, D.M. (2011) RAP80-directed tuning of BRCA1 homologous recombination function at ionizing radiation-induced nuclear foci. Genes Dev, 25, 685–700.

29. Li, X. and Manley, J.L. (2005) Inactivation of the SR protein splicing factor ASF/SF2 results in genomic instability. Cell, 122, 365–378.

