## Supplemental Figures for "DHX9-dependent recruitment of BRCA1 to RNA is required to promote DNA end resection in homologous recombination"

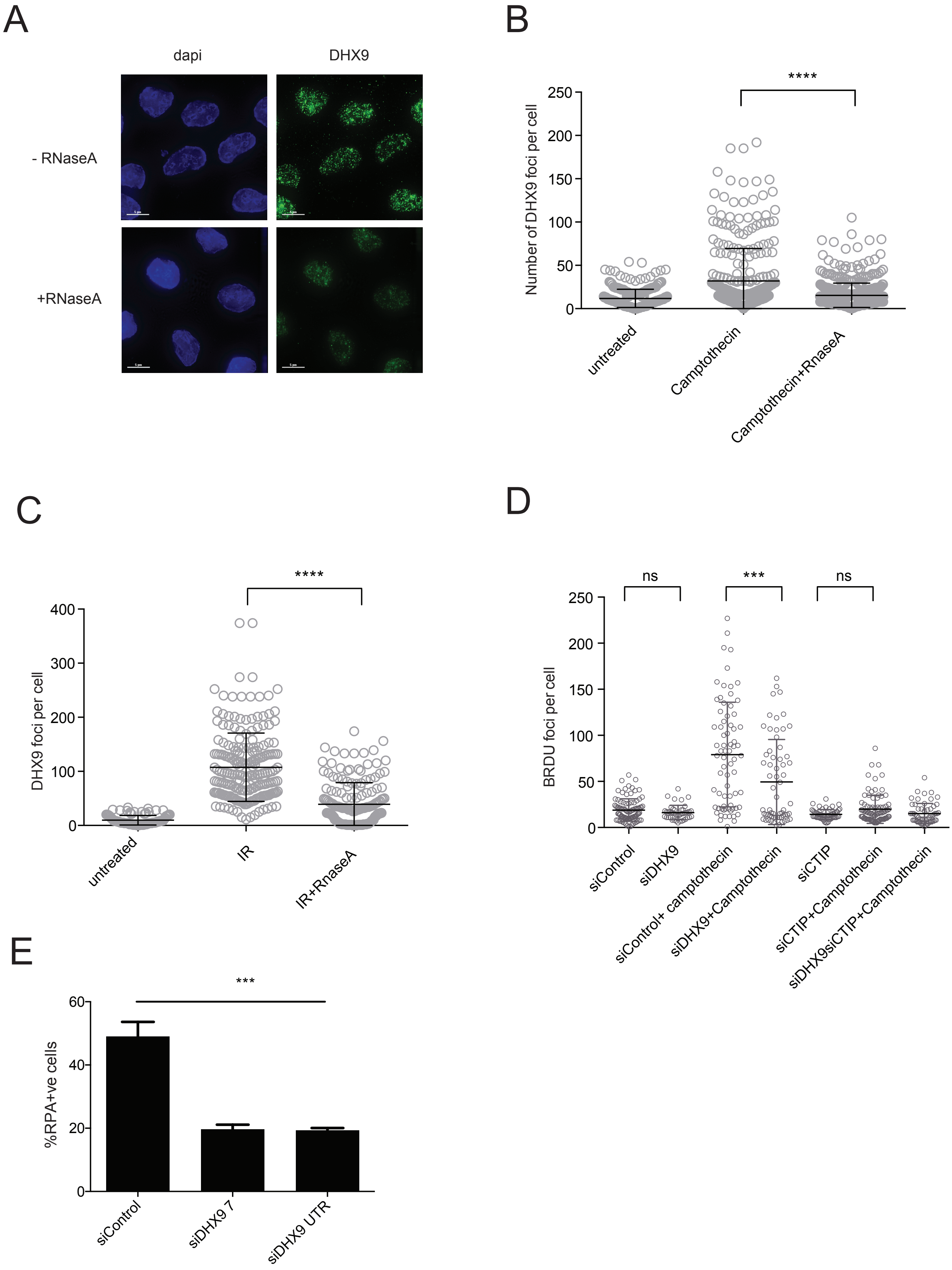


**Supplementary Figure 1.** (A) Fluorescent images showing that DHX9 nuclear foci formed after treatment of cells with camptothecin are disassembled by treating cells with RNaseA. B. Quantification of experiment in A. (C) DHX9 nuclear foci formed after treatment of cells with ionizing radiation are disassembled by treating cells with RNaseA. (D) Resection of DNA ends to form ssDNA is diminished in cells knocked down for DHX9 and CtIP. ssDNA was measured by staining for accessible BrdU. (E) DHX9 defective cells are impaired for recruitment of RPA to DNA damage. Percentage cells staining with RPA after treatment with camptothecin, measured using FACS. Statistical significance was determined (B-D) using Mann Whitney test (***$p \leq 0.001$, ****$p \leq 0.0001$) and (E) using one-way Anova

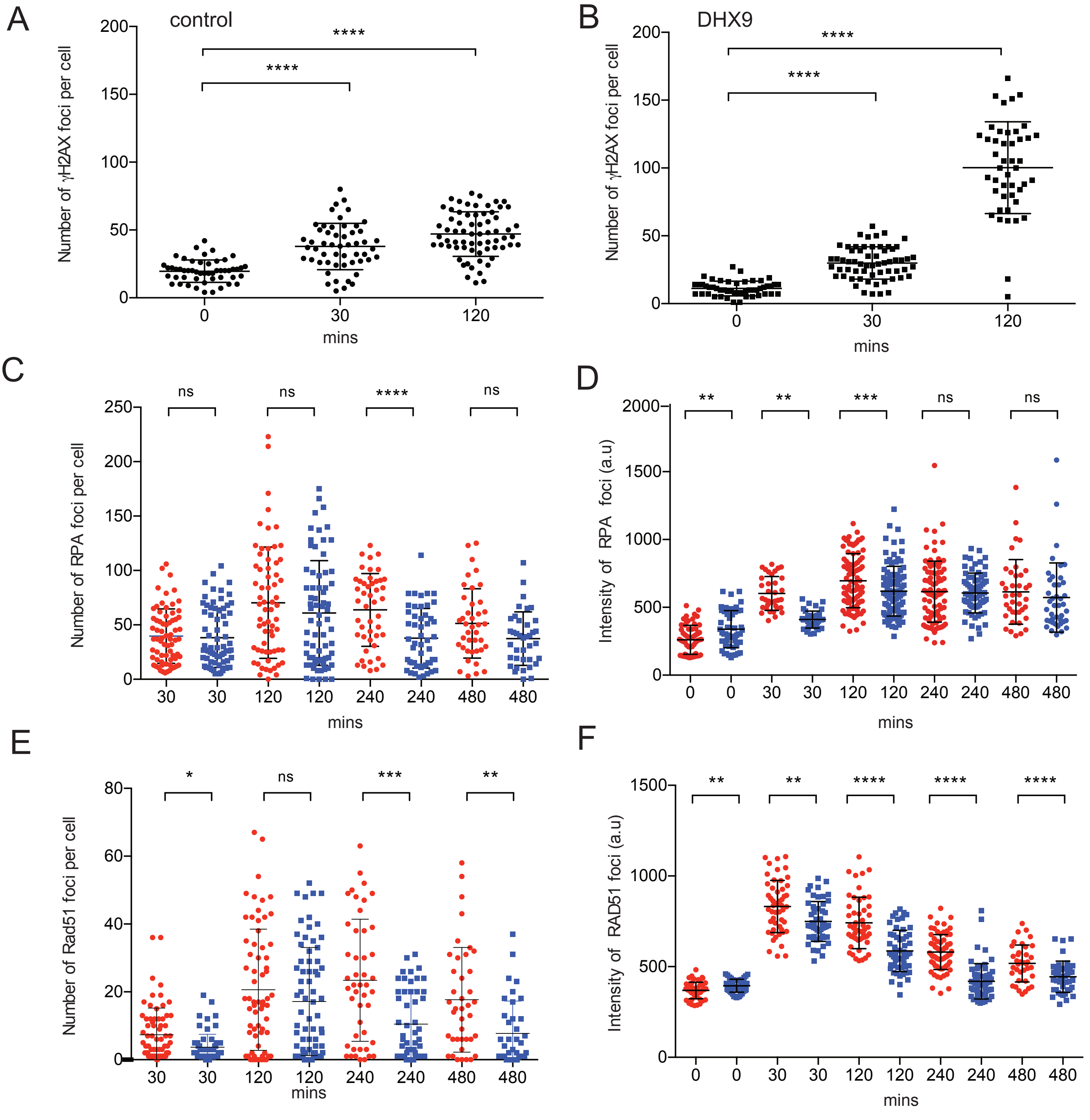


**Supplementary Figure 2.** (A) Number of γH2AX foci per cell after treatment with IR in siControl cells (B) Number of γH2AX foci per cell after treatment with IR insiDHX9 cells (C). Number of RPA foci per cell after treatment with IR in siControl (red) and siDHX9 cells (blue) (D). Intensity of nuclear RPA staining in cells treated with IR in siControl (red) and siDHX9 cells (blue). (E). Number of RAD51 foci per cell after treatment with IR in siControl (red) and siDHX9 cells (blue) (F). Intensity of nuclear RAD51 staining in cells treated with IR in siControl (red) and siDHX9 cells (blue). and siDHX9 cells. Statistical significance was determined using Mann Whitney test (*$p < 0.1$**,$p < 0.01$***,$p < 0.001$, ****$p < 0.0001$)
